# Zika virus infects pericytes in the choroid plexus and enters the central nervous system through the blood-cerebrospinal fluid barrier

**DOI:** 10.1101/841437

**Authors:** Jihye Kim, Michal Hetman, Eyas M. Hattab, Joshua Joiner, Brian Alejandro, Horst Schroten, Hiroshi Ishikawa, Dong-Hoon Chung

## Abstract

Zika virus (ZIKV) can infect and cause microcephaly and Zika-associated neurological complications in the developing fetal and adult brains. In terms of pathogenesis, a critical question is how ZIKV overcomes the barriers separating the brain from the circulation and gains access to the central nervous system (CNS). Despite the importance of ZIKV pathogenesis, the route ZIKV utilizes to cross CNS barriers remains unclear.

Here we show that in mouse models, ZIKV-infected cells initially appeared in the periventricular regions of the brain, including the choroid plexus and the meninges, prior to infection of the cortex. The appearance of ZIKV in cerebrospinal fluid (CSF) preceded infection of the brain parenchyma. We show that ZIKV infects pericytes in the choroid plexus, and that ZIKV infection of pericytes is dependent on AXL receptor tyrosine kinase. Using an in vitro Transwell system, we highlight the possibility of ZIKV to move from the blood side to CSF side, across the choroid plexus epithelial layers, via a nondestructive pathway (e.g., transcytosis). Finally, we demonstrate that brain infection is significantly attenuated by neutralization of the virus in the CSF, indicating that ZIKV in the CSF at the early stage of infection might be responsible for establishing a lethal infection of the brain. Taken together, our results suggest that ZIKV invades the host brain by exploiting the blood-CSF barrier rather than the blood-brain barrier.

**AUTHOR SUMMARY:** Zika virus invades the human brains and causes Zika-associated neurological complications; however, the mechanism(s) by which Zika virus accesses the central nerves system remain unclear. Understanding of the cellular and molecular mechanisms will shed light on development of novel therapeutic and prophylactic targets for Zika virus and other neurotropic viruses. Here we use in vivo and in vitro models to understand how Zika virus enters the brain. In mouse models, we found that Zika virus infects pericytes in the choroid plexus at very early stages of infection and neutralization of Zika virus in the cerebrospinal fluid significantly attenuate the brain infection. Further we show evidence that Zika virus can cross the epithelial cell layers in the choroid plexus from the blood side. Our research highlights that ZIKV invades the host brain by exploiting the blood-CSF barrier rather than the blood-brain barrier.

## INTRODUCTION

The recent Zika virus (ZIKV) outbreaks in the Americas clearly revealed that ZIKV can cross various biological barriers and establish infections in body compartments that are separated from the general circulation. ZIKV is sexually transmittable over genital barriers(1), and ZIKV is often found in the eyes, which are protected from the circulation by the blood–retinal barrier(2, 3). Importantly, ZIKV invades the brain of both fetuses and adults. Successful invasion of the central nervous system (CNS) by ZIKV causes detrimental ZIKV-associated neurological complications such as microcephaly(4, 5), Guillain-Barré syndrome(6), encephalitis, myelitis, and meningoencephalitis in infants and adults (7–12). In the context of infection of the CNS, which is separated from the circulation by a system of barriers, a critical question is how ZIKV gains access to the CNS from the circulation. Considering the fact that most nonneurological infections of ZIKV typically cause no to mild symptoms in humans, it is important to define the mechanism by which ZIKV crosses the barrier around the brain. Understanding of such mechanisms would shed light on novel therapeutic and prophylactic targets for ZIKV-induced neurological diseases.

The current speculation is that ZIKV crosses the BBB by productive infection of endothelial cells of the BBB(13, 14). It was previously shown that ZIKV infects primary human brain endothelial cells and is released from the basolateral side of the cell, indicating the potential release of virus from the endothelial cells into the cortex. However, no in vivo model studies have shown a clear understanding of how ZIKV invades the brain. Alternatively, an in vitro model using differentiated brain endothelial cells showed the potential of ZIKV to penetrate brain endothelial cell layers, which comprise the blood-brain barrier (BBB), without disrupting the structural integrity(15). The infection of astrocytes, which are in close contact with endothelial cells, is known to be associated with breakdown of the BBB(16), which could provide a route to the brain. However, it remains unclear how ZIKV initially infects astrocytes or how ZIKV penetrated brain endothelial layers.

To gain insight into the mechanism by which ZIKV invades the brain, we employed interferon-deficient mouse models and analyzed the early events of brain infection. Using the models, we show that ZIKV infection in the brain starts from the infection of pericytes in the choroid plexus followed by emergence of ZIKV in the cerebrospinal fluid (CSF). Furthermore, using an in vitro primary human brain vascular pericyte model, we show that AXL is a key factor for the productive infection of pericytes by ZIKV. ZIKV crossed choroid plexus epithelial layers via a nondestructive pathway (e.g., transcytosis) and that ZIKV transport across the choroid epithelial layers could be prompted by cellular factors from ZIKV-infected pericytes in a Transwell system of transformed human choroid plexus epithelial cells and human brain vascular pericytes. We demonstrate that the ZIKV circulating in the CSF at the early stage of infection might be the source of the infection of the cortex by employing an intrathecal administration of neutralizing antibody prior to infection of the brain. Our data collectively suggest that ZIKV might invade the host brain by infection of pericytes followed by transcytosis across the epithelial layers in the choroid plexus, the blood-CSF barrier.

## RESULTS

### ZIKV Infection of the choroid plexus and the meninges preceded infection of the cortex in mouse models

Several studies using type-1 interferon receptor knockout mice have shown that when ZIKV is administered via subcutaneous, intraperitoneal, or intravenous routes, it establishes a robust brain infection in adult mice(17–19). To understand the mechanism by which ZIKV gains access to the brain, we investigated an early stage of brain infection in type-1 interferon receptor knockout mice, B6(Cg)-Ifnar1tm1.2Ees/J (hereafter referred to as IFNAR^-/-^) with the PLCal_ZV strain of ZIKV. PLCal_ZV is an early Asian lineage Zika virus with an alanine-to-valine amino acid substitution at residue 188 in NS1(20) (GenBank KF993678.1), which leads to higher NS1 antigenemia and infectivity in the host compared to those of earlier isolates such as the FSS13025 strain(21, 22). IFNAR^-/-^ mice were infected with ZIKV subcutaneously, and the brains were harvested at 2, 3, and 4 days post infection (DPI). The brains were analyzed with in situ chromogenic RNA hybridization (hereafter, RNAScope assay) with a specific probe against the ZIKV genome. This method provided specific detection of ZIKV RNA in tissue mounted on slides. In our model, ZIKV-positive cells first appeared at 3 DPI in the choroid plexus (CP) and the meninges in the mouse brains (Fig. 1). While particle-like ZIKV RNA stains were also detected within the brain capillaries (Fig. 1 bi, gray arrowhead), no infected cells were detected in the capillaries of the cortex at 3 DPI. The CPs in all ventricles (i.e., lateral, third, and fourth ventricles) and the majority of meninges consistently showed strong positive signals for ZIKV in all samples (4 brains per timepoint) at 3 and 4 DPI (Fig. 1 bii and cii, **black arrowhead or black arrow**). In the cortex, ZIKV-infected cells appeared for the first time at 4 DPI. At the early time points, the infection pattern in the cortex was focal and not evenly distributed throughout the entire brain area (Fig. 1c, **black arrowheads)**. This infection pattern was remarkably different from the brains infected with Venezuelan equine encephalitis virus (strain TC-83 in AG129 mice, Fig. 1d **and** Supplementary Fig. 1) that showed widely distributed strong positive staining around the capillaries in the cortex as previously reported by others(23).

**Fig. 1.**
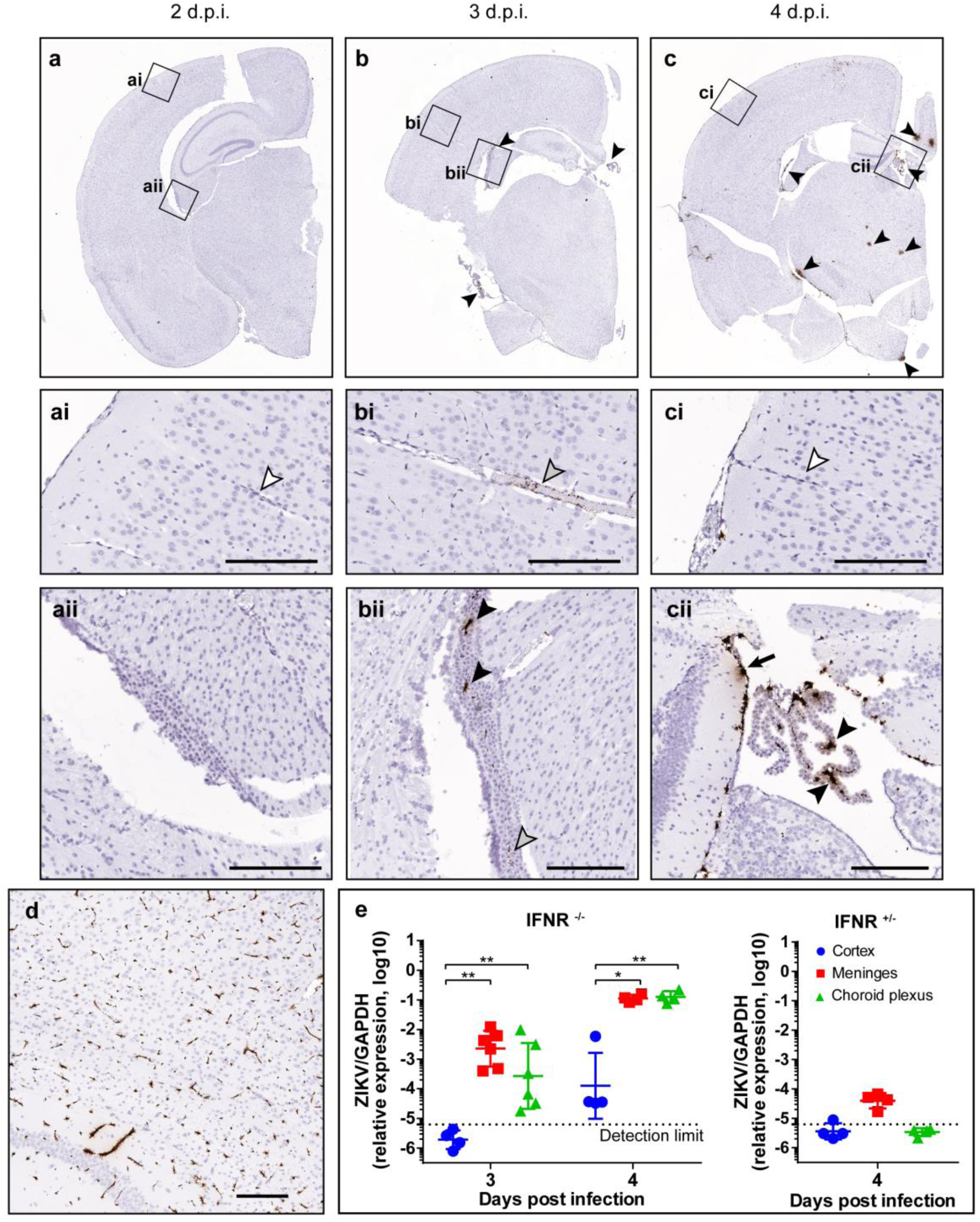
ZIKV infection of the brain in mice starts from the infection of the choroid plexuses and meninges. **a-c**, IFNAR^-/-^ mice (n=4 per time point) were subcutaneously infected with 1000 p.f.u. of ZIKV and euthanized at 2, 3, and 4 DPI. The brains were subjected to an in-situ RNA hybridization (RNAScope) assay with a probe detecting ZIKV RNAs. Positive signals for the target RNA are shown in dark brown, whereas the counterstaining with hematoxylin is shown in light blue. Images taken with a 20X objective were stitched with Hugin panorama image stitch software. Black, gray, and white arrowheads indicate virus-infected cells, virus, and the absence of ZIKV-specific staining, respectively. Detailed images of the areas outlined with a black box are shown in ai, bi, and ci (cortex and capillaries) and aii, bii, and cii (choroid plexus). Scale bars = 200 µm. **d**. AG129 mice (n=3 per group, two experiments) were subcutaneously infected with 1000 p.f.u. of TC-83 and euthanized at 2 days post infection. The TC-83 vRNA was visualized in an RNAScope assay with a VEEV-specific probe. Scale bars = 200 µm. **e**, IFNAR^-/-^ mice (left, n=4–6/group) or IFNAR^+/-^ mice (right, the cage mate n=4) were infected with 1000 p.f.u. of ZIKV subcutaneously, and the brains were harvested at 3 and 4 DPI. The viral loads in the cortex (blue circles), meninges (red squares), and choroid plexuses (green triangles) were determined with a ZIKV-specific qRT-PCR assay and normalized to the expression of GAPDH mRNA in each sample. Data were analyzed by one-way ANOVA with Tukey’s multiple comparison tests. * *P* = 0.003, ** *P* =0.0001.

To confirm that the infection in the choroid plexus and meninges preceded that in the cortex, we compared the viral loads in these tissues using the quantitative real-time PCR (qRT-PCR) method (Fig. 1e). IFNAR^-/-^ mice infected with ZIKV as above were euthanized at 3 and 4 DPI and subjected to cardiac perfusion prior to the harvest of the brain to remove blood in the neurovascular systems. At 3 DPI, significant amounts of viral RNA were detected in the meninges and the CP but not in the cortex of the same brains. In the cortex, ZIKV appeared at 4 DPI. This result confirmed our findings with the chromogenic in situ RNA hybridization assay. The negative control group, cage mate IFNAR^+/-^ mice infected with ZIKV, did not show any measurable amounts of viral RNA in all three tissues, confirming that spread to the brain might require IFNAR1 deficiency(16).

To understand if the infection of CP and meninges prior to the infection of the cortex is unique to this strain of ZIKV and mouse, we employed another mouse model, the AG129 model (IFN-α/β/γ receptor knockout strain), and other ZIKV isolates (DAK AR 41524 and PRVABC59) representing African and American lineages in addition to PLCal_ZV(4, 18). We found that the progress of ZIKV infection in the brains of AG129 mice was nearly identical to that of IFNAR^-/-^ mice. For the PLCal_ZV strain of ZIKV infection in AG129 mice, ZIKV-infected cells were found in the CP and the meninges at 4 DPI, and very little viral RNA staining was detected within the cerebral cortex region (Supplementary Fig. 2). Robust infection in the cortex was observed at 6 DPI. Again, little or no infection of the capillaries was found. These same findings – earlier infection of the meninges and CP than of the cortex – were observed for the DAK AR 41524 and PRVABC59 strains, indicating that infection of the CP and meninges is a common feature of ZIKV in these mouse models (Supplementary Fig. 3).

**Fig. 2.**
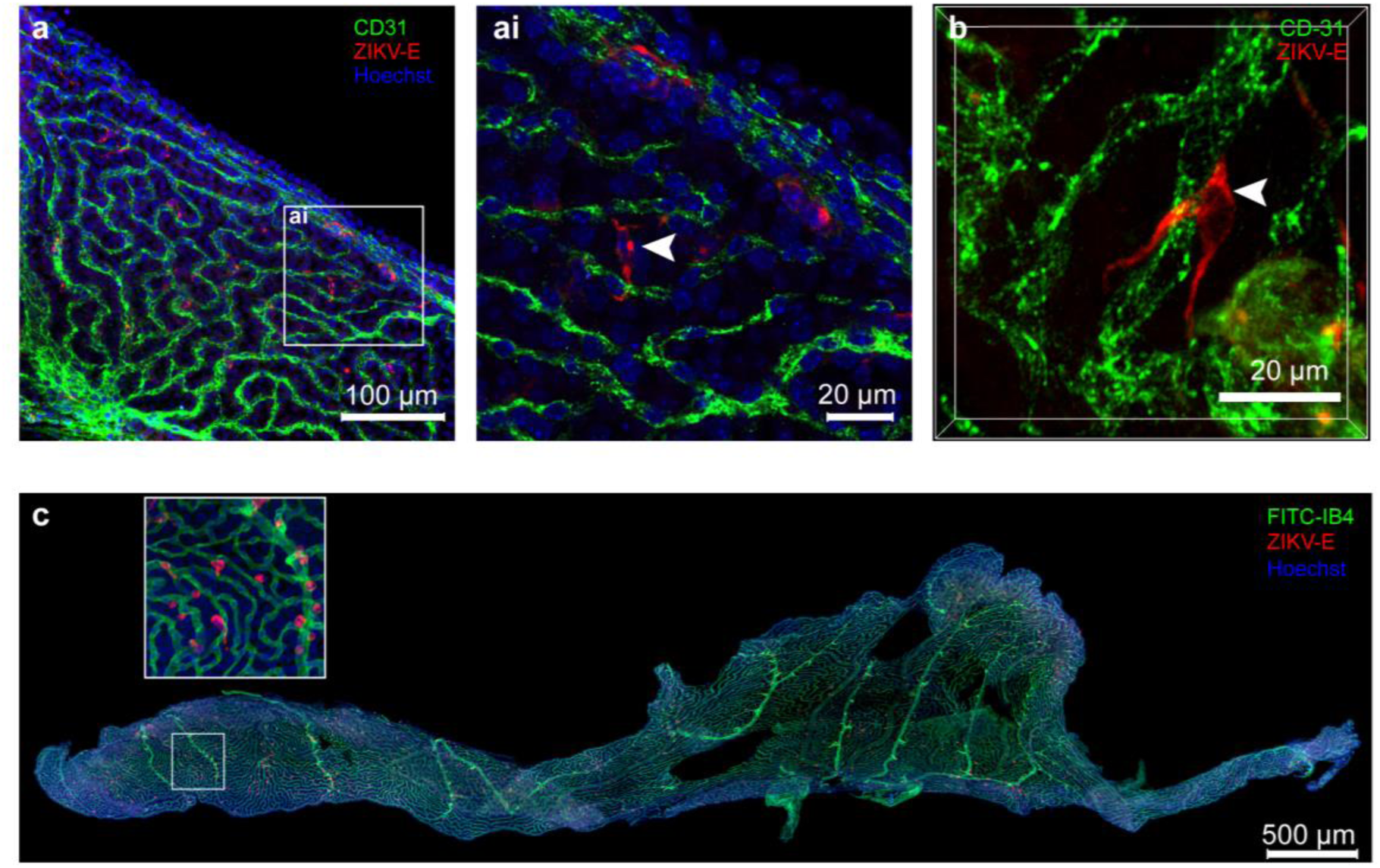
ZIKV-infected cells in the choroid plexus are tightly associated with endothelial cells. **a,** The lateral choroid plexuses from ZIKV-infected IFNAR^-/-^ mice (n=4 per time point) were harvested at 4 DPI and subjected to whole mount IFA assay. The whole tissue was stained with anti-CD31 antibody (endothelial cells, green), hu-4G2 (ZIKV envelope, red), and Hoechst 33342 (nuclei, blue) and then mounted for evaluation by confocal microscopy Scale bar = 100 µm. **ai**, A higher resolution image of the inset area of Fig 3 a. Scale bars = 20 µm. **b**, A representative image of ZIKV-infected cells tightly connected with endothelial cells. Scale bars = 20 µm. **c**, The lateral choroid plexus of AG129 mice infected with ZIKV (1000 p.f.u. at 4 DPI) were stained with FITC-IB4 (endothelial layer, green), hu-4G2 (ZIKV envelope, red), and Hoechst 33342 (nuclei, blue). A total of 74 images taken with a 20X objective were stitched together with Hugin image stitch software. Scale bar = 500 µm.

**Fig. 3.**
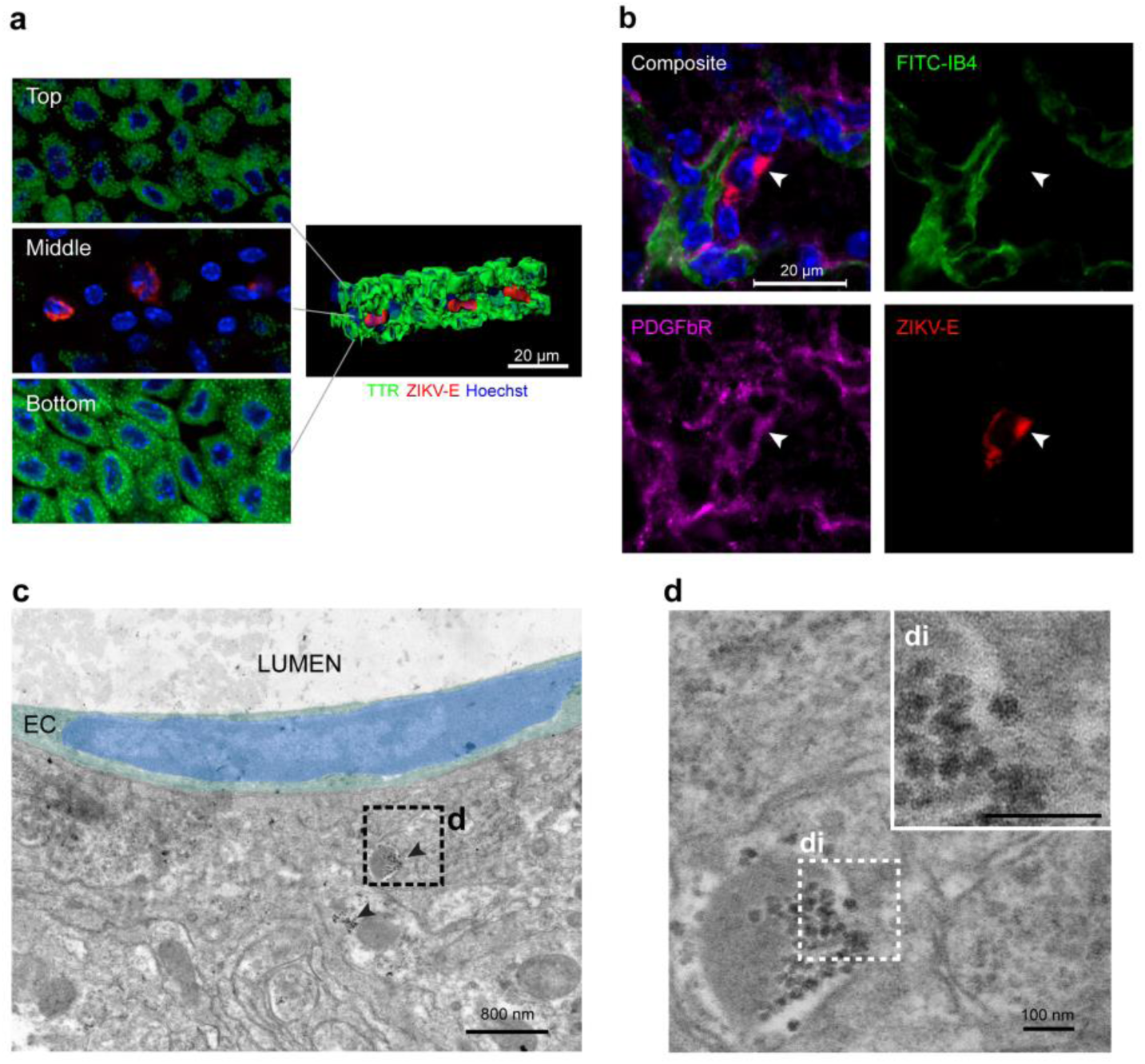
ZIKV infects pericytes in the choroid plexus. **a-d**, IFNAR^-/-^ mice were infected with ZIKV as above, and the choroid plexuses were isolated at 4 DPI. **a**, Choroid plexuses were stained with antibodies against TTR (choroid plexus epithelial cell maker, green) and ZIKV-E protein (red). **left**: three layers of confocal images (labeled with Top, Middle, and Bottom) along the Z-axis. **right**: a three-dimensional reconstruction of the z-stacked image, highlighting the ZIKV-infected cells in the stroma layer of the choroid plexus. The top layer, middle, and bottom layers showed TTR+/ZIKV-, TTR-/ZIKV+, and TTR+/ZIKV-, respectively. A representative image is shown. **b**, Choroid plexuses were costained with FITC-IB4 for capillaries (green), anti-PDGFβR antibody for pericytes (purple), and anti-ZIKV-E antibody for the Zika envelope protein (red). Nuclei were stained with Hoechst 3342 stain (blue). The ZIKV-E signal colocalized with PDGFβR. For **a** and **b**, scale bars = 20 µm. **c**, An electron micrograph showing a pericyte infected with ZIKV on a choroid plexus capillary. Pseudocolors (green and blue) indicate the cytoplasm and the nucleus, respectively, of a choroid plexus endothelial cell (EC) facing the lumen of a capillary (LUMEN). Patches of electron-dense particles (arrowheads) were found in a pericyte next to the EC. **d**, A higher magnification of the image from **c** shows that the particles have characteristics typical of flavivirus nucleocapsids. **di**, A higher magnification image of the inset area of Fig **d**, Scale bars: 800 nm in **c** and 100 nm in **d and di**.

Overall, these experiments revealed that ZIKV infects the choroid plexus and meninges in the brain, and the infection in these tissues precedes the infection of other regions in the brain.

### ZIKV infects pericytes, not endothelial cells in the CP

The strong ZIKV RNA signal in the CP and the unique biology of the CP led us to further investigate the CP as the ZIKV target tissue for brain invasion. The CP is a highly vascularized, specialized tissue located in the brain ventricles. The choroid plexus consists of a central stroma covered by two epithelial layers: the polarized choroid plexus epithelium facing the stroma at the basolateral side and the CSF at the apical side. Primary function of the CP is the production of CSF by transporting water and biomolecules between the blood and the CSF(5, 24). The production of CSF by the CP is mainly achieved by three cell types: 1) fenestrated endothelial cells that form capillaries, 2) pericytes, which are in direct contact with the endothelial cells and are responsible for regulating permeability, cerebrospinal blood flow, angiogenesis, clearance of debris, neuroinflammation, and stem cell activity(25–27), and 3) CP epithelial cells, which serve as the selective barrier between the blood and the CSF(24,28,29). The tight junctions between the CP epithelial cells limit the paracellular diffusion of molecules across the barrier, and the transport of molecules is selectively achieved by transcytosis in CP epithelial cells(30, 31). Interestingly, unlike the blood vessels that comprise the blood-brain barrier in the parenchyma, the blood vessels in the choroid plexus are highly fenestrated and leaky, which can be an ideal milieu for infectious agents to easily spread into the tissue from the circulation system(32, 33).

We sought to define the cell type that initiates ZIKV brain infection in the CP. To understand the characteristics of ZIKV-infected cells in the CP, we employed confocal microscopy of whole-mount CP tissues. Consistent with the time course histochemistry results (Fig. 1), we observed that ZIKV envelope(E)-positive cells were found in the choroid plexus (Fig. 2). No endothelial cells (CD31+) showed positive staining for ZIKV; rather, ZIKV-infected cells were always in close contact with endothelial cells. High-resolution images showed that ZIKV-positive signals were always found in perivascular cells attached to capillaries or connecting two capillaries **(as shown in** Fig. 2ai and b, **white arrowhead).** A three-dimensional reconstitution analysis showed that the ZIKV-infected cells are tightly associated with the endothelial cells (Supplementary Fig. 4). We confirmed this finding using secondary reagents: FITC-labeled *Griffonia simplicifolia* (Bandeiraea) isolectin B4 (IB4) and CPs isolated from ZIKV-infected AG129. IB4 is a lectin that can bind the basement membrane of capillaries or macrophages(34). The CPs from ZIKV-infected AG129 mice also showed ZIKV E-positive cells located near endothelial cells throughout the entire CP (Fig. 2d), which confirmed that ZIKV-positive cells were not endothelial cells but were in proximity of the capillaries in the CP.

**Fig. 4.**
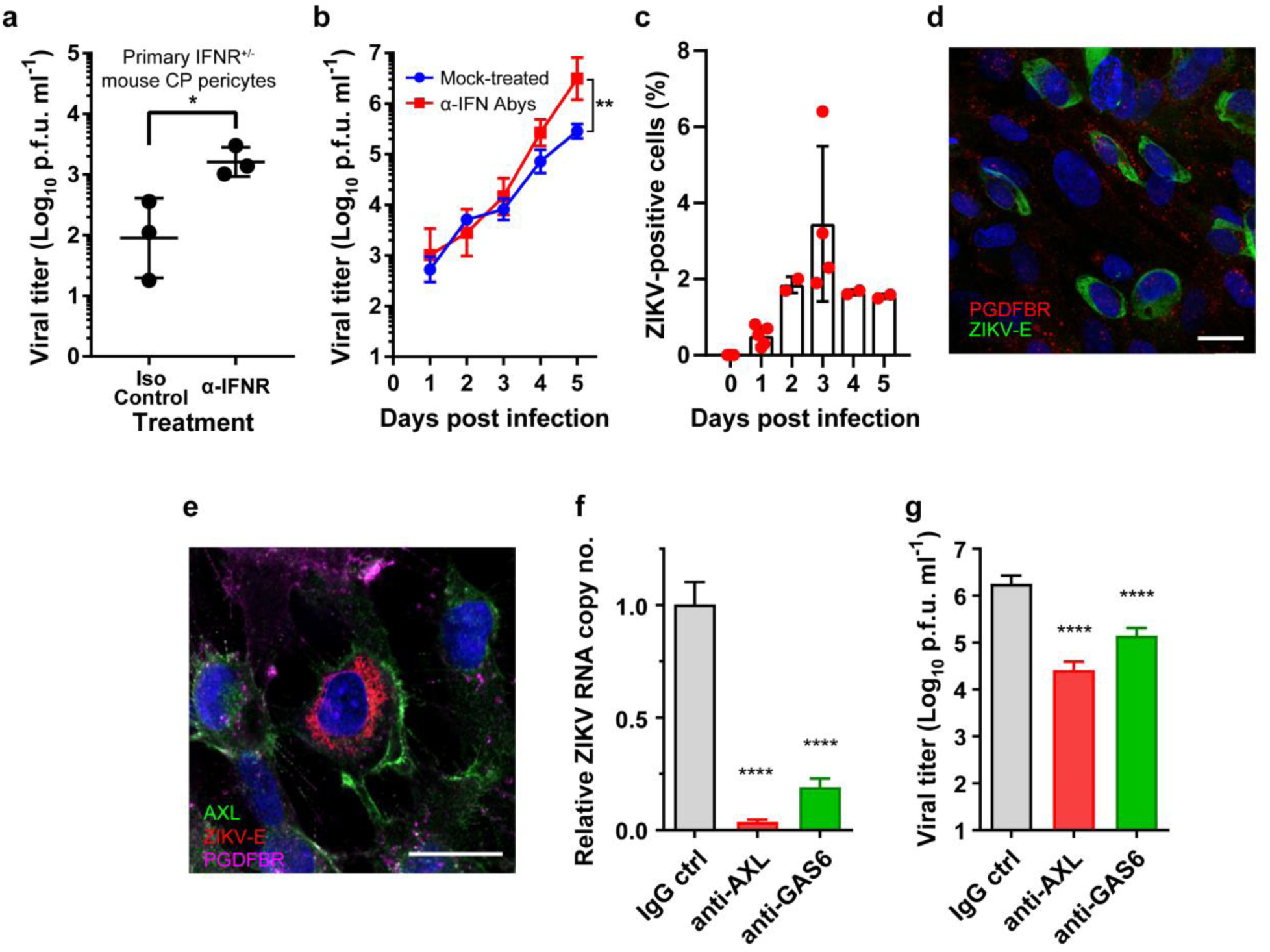
ZIKV infects human brain vascular pericytes in an AXL-dependent manner. **a**, ZIKV replication in primary pericytes from mouse choroid plexus (IFNAR^+/-^). Cells were infected with ZIKV, PLCal_ZV (MOI = 0.1) and grown in the presence of 5 µg/mL mouse anti-IFNAR-1 neutralizing antibody (clone MAR 5A3, ɑ-IFNAR) or isotype control antibody (clone MOPC-21). The cell culture supernatant was harvested at 5 DPI, and the virus titer was enumerated. **b**, Primary human brain vascular pericytes were infected with ZIKV as in **a** and then incubated in the presence/absence of a human type 1 IFN neutralizing antibody mixture (ɑ-IFN Aby). The supernatants were harvested every 24 hours for five days, and the virus titers were enumerated with a virus titration assay. * P <0.05 (Student’s t-test) and ** P < 0.005 (two-way ANOVA). **c**, Primary human brain vascular pericytes were infected with ZIKV and ZIKV-infected cells were enumerated by using FACS with anti ZIKV-E (clone 4G2). **d**, HBVP infected with ZIKV were stained with antibodies against PDGFβR (red) and ZIKV-E protein (green). Two consecutive confocal layers were projected into a single layer with the Z-project function of ImageJ software (version 2.0.0). **e**, HBVP infected with ZIKV were stained with antibodies against AXL (green), PDGFβR (magenta) and ZIKV-E protein (red). Scale bars = 20 µm. **f** and **g**, HBVP were pre-treated with antibody for three hours then infected with ZIKV (m.o.i. = 1). Three days later, viral RNA (**f**) and progeny virus titers in the supernatants (**g**) were analyzed. **** P < 0.001 (one-way ANOVA).

Furthermore, we sought to determine the specific cell type that was infected by ZIKV in the CP. First, we confirmed that ZIKV-positive cells are not located in the CP epithelial cell layers (Fig. 3a). CP epithelial cells are connected to each other with tight junction proteins (e.g., ZO-1) at the apical and basolateral sides of the CP. Immunofluorescence microscopy using anti-TTR (i.e., transthyretin, a CP epithelial cell marker) antibody(35, 36) showed that ZIKV-positive cells were located in the CP stroma space between the CP epithelial layers, not within the epithelial layers. This finding is different from what has been shown in a mouse model of Chikungunya virus infection, where the CP epithelial cells were infected(37).

Based on the morphology and the location of the ZIKV-infected cells within the CP (Fig. 3a), we hypothesized that the ZIKV-infected cells are pericytes. Pericytes are in direct contact with endothelial cells and continuously communicate with endothelial and other cells via secretory cytokines(38, 39). Brain vascular pericytes are responsible for regulating BBB permeability, cerebrospinal blood flow, angiogenesis, clearance of debris, neuroinflammation, and stem cell activity(25–27). To test this hypothesis, we employed confocal fluorescence microscopy of whole-mount CPs (Fig. 3b) with an antibody against platelet-derived growth factor receptor beta (PDGFβR), a marker for pericytes (26,40,41). The ZIKV-positive cells in the CP showed positive staining for PDGFβR and were found on the outer side of capillaries, indicating that the ZIKV-infected cells were indeed pericytes. We also confirmed the replication of ZIKV in CP pericytes by using transmission electron microscopy (TEM) of ZIKV-infected CP (Fig. 3c and 3d). Electron-dense particles with morphological characteristics typical of the nucleocapsid of flaviviruses were found on the surface of a vesicle in a cell adjacent to an endothelial cell on a CP capillary. The size of the particles was 28.2 ± 2.2 nm (mean and standard deviation from measurements of 28 particles), which is consistent with the size of immature ZIKV nucleocapsid cores previously determined by others(42). Our results clearly showed that ZIKV targets pericytes not endothelial or epithelial cells in the CP.

### Pericytes from mouse CP and human brains are susceptible to ZIKV infection

Next, we sought to model ZIKV infection of pericytes in vitro and examined the IFN dependence of ZIKV infection of pericytes. To test these models, we infected primary cultured mouse CP pericytes isolated from IFNAR^+/-^ mice (Fig. 4a) and primary human brain vascular pericytes (HBVPs) (Fig. 4b) with ZIKV in the presence or absence of type I IFN neutralizing antibodies (NAbs). The titers of progeny viruses produced from the cells were determined as proof of viral replication in the cells. We found that ZIKV was able to replicate and produce progeny virus in both cell types, indicating that pericytes are indeed susceptible to ZIKV infection. The mouse CP pericytes produced a low titer of progeny viruses in the absence of NAb against mouse type 1 interferon receptor (IFNAR-1). A slightly but significantly increased amount of progeny viruses was generated in the presence of NAb against mouse IFNAR-1 (mean virus titers at 5 DPI of 162 and 1796 p.f.u./mL for isotype control and anti-IFNAR Ab, respectively, Fig. 4a). Because there are no commercially available human choroid plexus pericytes, we employed HBVP cells as our human pericyte model. HBVPs were more susceptible to ZIKV infection than mouse CP pericytes (Fig. 4b), resulting in an increase in virus titer during the infection with a significant virus titer (mean titer of 2.93 x 10^5^ p.f.u./mL at 5 DPI). ZIKV-infected HBVPs grown in the presence of NAbs against human type 1 IFNs and IFNAR-1 produced an even higher amount of progeny virus (mean virus titer of 4.11 x 10^6^ p.f.u./mL). Interestingly, not all HBVPs were susceptible to ZIKV. Only small fractions of HBVPs, approximately 1 to 3.5% compared to the total number of cells at its peak at 3 DPI, were positive for ZIKV (Fig. 4c). These ZIKV-positive cells were positive for PDGFβR, a pericytes marker (Fig. 4d). Our results clearly showed that mouse CP pericytes and human brain pericytes are susceptible to ZIKV infection and that the type 1 IFN system plays a role in controlling ZIKV replication in these cells.

### AXL is a key factor for a productive infection of ZIKV in brain pericytes

TAM (TYRO3, AXL, MER) receptor tyrosine kinases are known important host factors that determine the susceptibility to ZIKV as a cellular receptor for viral entry and/or an attenuator of innate antiviral response(43). Notably, AXL receptor tyrosine kinase (AXL) is known to be important for efficient infection of human endothelial cells, skin cells, Sertoli cells, astrocytes, and brain glial cells (44–47). However, some evidence indicates that the dependency of ZIKV on TAM receptors for infection might be cell-type or model specific (e.g., IFN dependency) (48, 49). Here we assessed if the AXL protein plays role in the productive infection of pericytes. First, using qRT-PCR, we detected a significant AXL expression level in HBVP (Supplementary Fig. 5). We also confirmed that ZIKV-infected HBVP (i.e., ZIKV E and PDGFBR-positive) stained positively for AXL (Fig. 4e). Next, to understand the functional role of AXL in ZIKV infection, we examined the effect of blocking of AXL and growth arrest specific gene-6 (GAS6), ligand for AXL, on ZIKV replication in HBVP. A pretreatment of HBVP with anti-AXL or anti-GAS6 antibodies resulted in a greater than 96% or 80% reduction, respectively, in viral replication measured by qRT-PCR (Fig. 4f) of intracellular viral RNA, as well as a reduction in progeny viral titers (Fig. 4g). Overall, our data clearly indicate that ZIKV-infection of HBVP is dependent on AXL and GAS6.

**Fig. 5.**
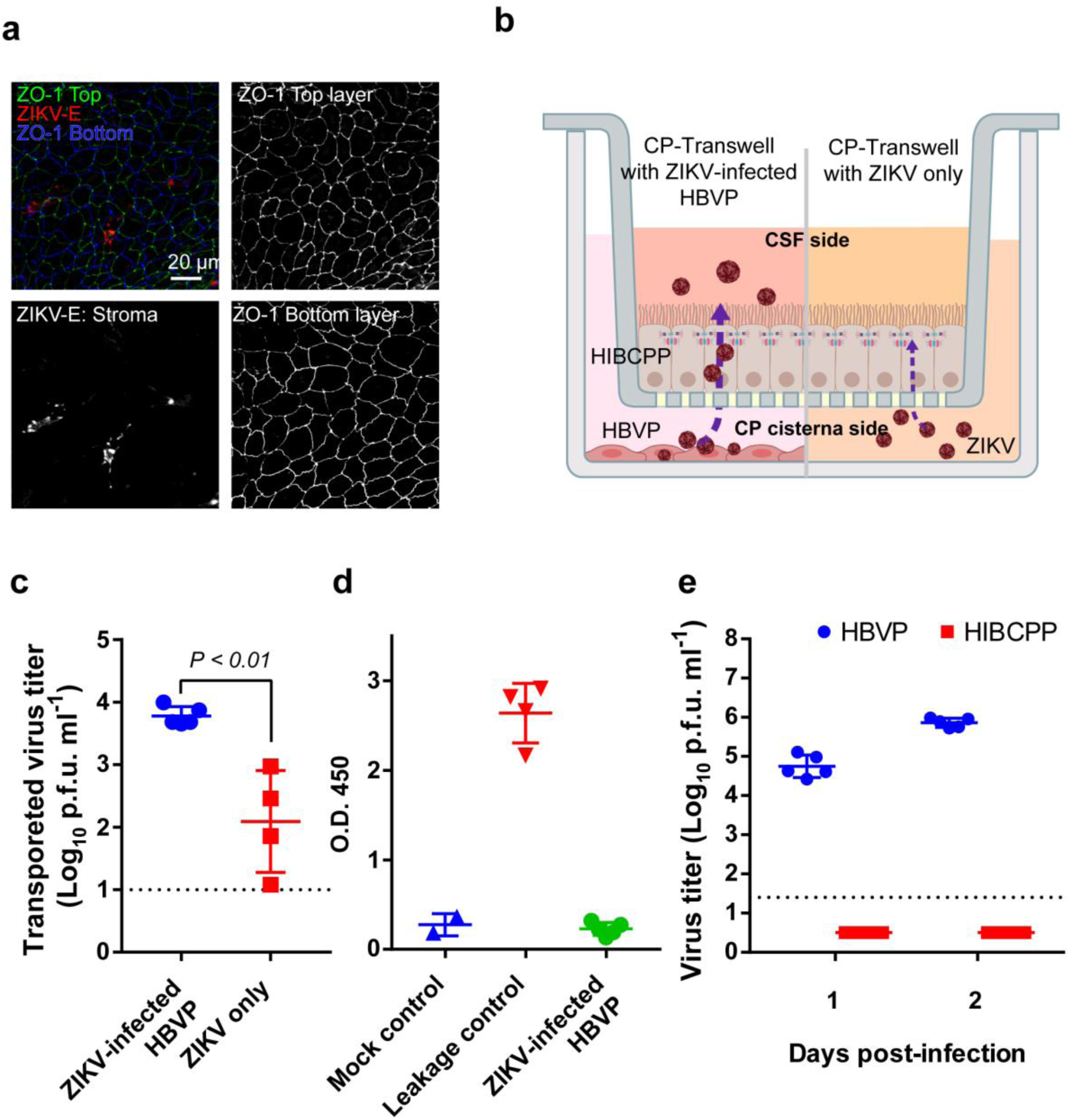
ZIKV crosses CP epithelial cell barriers without destruction of the tight junctions. **a**, CPs isolated from ZIKV-infected mice were subjected to immunostaining in a whole-mount choroid plexus assay with antibodies against ZO-1 and ZIKV-E. The entire depth of the area shown was imaged with a Zeiss LSM 710 Duo/LIVE 5 confocal microscope along the Z-axis, and the two epithelial cell layers (top and bottom) and stroma cell layers were extracted and projected into single layers with the Z-project function of ImageJ software (version 2.0.0). **b**, HIBCPP cells are grown and differentiated into a monolayer in Transwell inserts. Fully developed HIBCPP Transwell inserts (determined by TEER) are placed in wells with ZIKV-infected primary human brain pericytes grown in the lower chamber for two days. Virus transported to the apical side (CSF side) is collected and is quantitated by using a standard plaque assay or ZIKV-specific qRT-PCR. **c**. Fully developed HIBCPP barriers grown in Transwell inserts (n=4 or 5 per group) were basolaterally exposed to ZIKV-infected HBVP cells (2 DPI) or ZIKV diluted in HBVP-conditioned media. Twenty-four hours later, cell supernatants from the apical side were harvested, and the virus titers were determined. **d**, After 5 days of coculture with ZIKV-infected HBVP cells, the Transwell inserts with HIBCPPs were basolaterally exposed to HRPO-conjugated IgG for two hours, and the cell culture media on the apical sides were tested for HRPO activity. As a leakage control, a scratch was made on the HIBCPP cells fully grown in Transwell inserts. **e**, HBVP (blue circles) and HIBCPP (red squares) cells grown on 12-well plates were infected with ZIKV (m.o.i. = 0.1) and then incubated. The virus titers in the supernatants were enumerated with a virus titration assay.

### An in vitro blood-cerebrospinal fluid (B-CSF) barrier model recapitulates ZIKV entry into the CSF side

Then, we tested whether the tight junctions between CP epithelial cells were breached by the infection of the pericytes, which would allow virus to cross the B-CSF barrier and enter into the CSF. First, we employed a confocal microscopy assay for ZO-1 (zonula occluden-1, a tight junction protein) to observe changes in tight junctions in whole-mount CP tissue from ZIKV-infected mice. We found that tight junctions are rather intact; no destructions of ZO-1 junctions were identified in the ZIKV-infected CP (Fig. 5a).

To recapitulate the entry of ZIKV into the CSF through the B-CSF barrier, we developed an in vitro model utilizing the human choroid plexus papilloma (HIBCPP) cell line(50) and HBVP cells. HIBCPP is an established cell line derived from human CP epithelial cells and is known to form cellular barriers when it is grown and differentiated on Transwell membranes. HIBCPP cells grown and fully differentiated on Transwell membranes were exposed to ZIKV basolaterally by being placed in cell culture wells with HBVP cells that were infected with ZIKV for two days. After twenty-four hours of exposure, the amount of virus that crossed the HIBCPP cell barrier was determined by titration of virus in the cell culture supernatant in the apical side of the Transwells (Fig. 5b).

We found that a significant amount of ZIKV was transported to the apical chamber when the HIBCPP cell barrier was exposed to ZIKV-infected HBVP culture supernatant (Fig. 5c). The transported virus (a mean virus titer of 6.0 x 10^3^ p.f.u./mL, n=5) was approximately 6% compared to the amount of virus in the bottom chamber (mean virus titer of 1.0 x 10^5^ p.f.u./mL, n=5).

To understand whether active replication of ZIKV in pericytes plays a role in ZIKV crossing the B-CSF barrier, we also compared ZIKV directly diluted in media to ZIKV-infected HBVPs as the source of virus. HIBCPP cells grown in Transwells were basolaterally exposed to ZIKV diluted in conditioned media from uninfected HBVPs (1.0 x 10^5^ p.f.u./mL) for twenty-four hours, and the amount of virus in the apical chamber was determined.

We found that much less ZIKV was transported (a mean virus titer of 1.2 x 10^2^ p.f.u./mL, n=4) when ZIKV diluted in HBVP-conditioned media was used compared to that when cells were cocultured with ZIKV-infected HBVPs. This implies that cytokines or other cellular factors released from infected pericytes may play a role in promoting ZIKV to cross the B-CSF barrier. The integrity of the barrier was maintained after exposure to either ZIKV-infected HBVP cells or free ZIKV in HBVP-conditioned media (Fig. 5d).

HIBCPP cells were resistant to ZIKV infection, which indicates that virus found in the apical chamber was not progeny virus produced in HIBCPP cells (Fig. 5e). Our results on the lack of active replication in CP epithelial cells and preservation of the integrity of tight junctions in vitro and in vivo, suggest a nondestructive transport pathway (e.g., transcytosis) as the ZIKV brain entry mechanism.

### ZIKV present in the CSF at the early stage is important for brain infection

Our results so far implied that ZIKV might exploit the B-CSF barrier as the gateway to the brain in this mouse model. To test whether ZIKV exploits the B-CSF barrier to enter into the CSF, we first examined the presence of virus in the CSF prior to infection of the cortex. After IFNAR ^-/-^ mice were infected with ZIKV subcutaneously, the CSF and serum were collected at 2, 3, and 4 DPI, and the viral loads were determined (Fig. 6a). At 2 DPI, ZIKV was highly abundant in the serum with a median viral titer of 5.94 x 10^4^ p.f.u./mL; however, no ZIKV was found in the CSF of the same animals. A significant level of ZIKV (median virus titer of 5.43 x 10^3^ p.f.u./mL) was detected in the CSF at 3 DPI, at which time the CP was infected, but the cortex was not (Fig. 1). While the viral load in the serum decreased at 4 DPI (1.56 x 10^4^ p.f.u./mL) compared to 3 DPI (5.12 x 10^5^ p.f.u./mL), the viral load in the CSF continued to increase to a titer of 1.11 x 10^4^ p.f.u./mL at 4 DPI, which might be partially due to infection of the CP and cortex. This experiment clearly showed that the presence of ZIKV in the CSF precedes the infection of the cortex, indicating that ZIKV in the CSF might have directly originated from the blood across the B-CSF barrier.

**Fig. 6.**
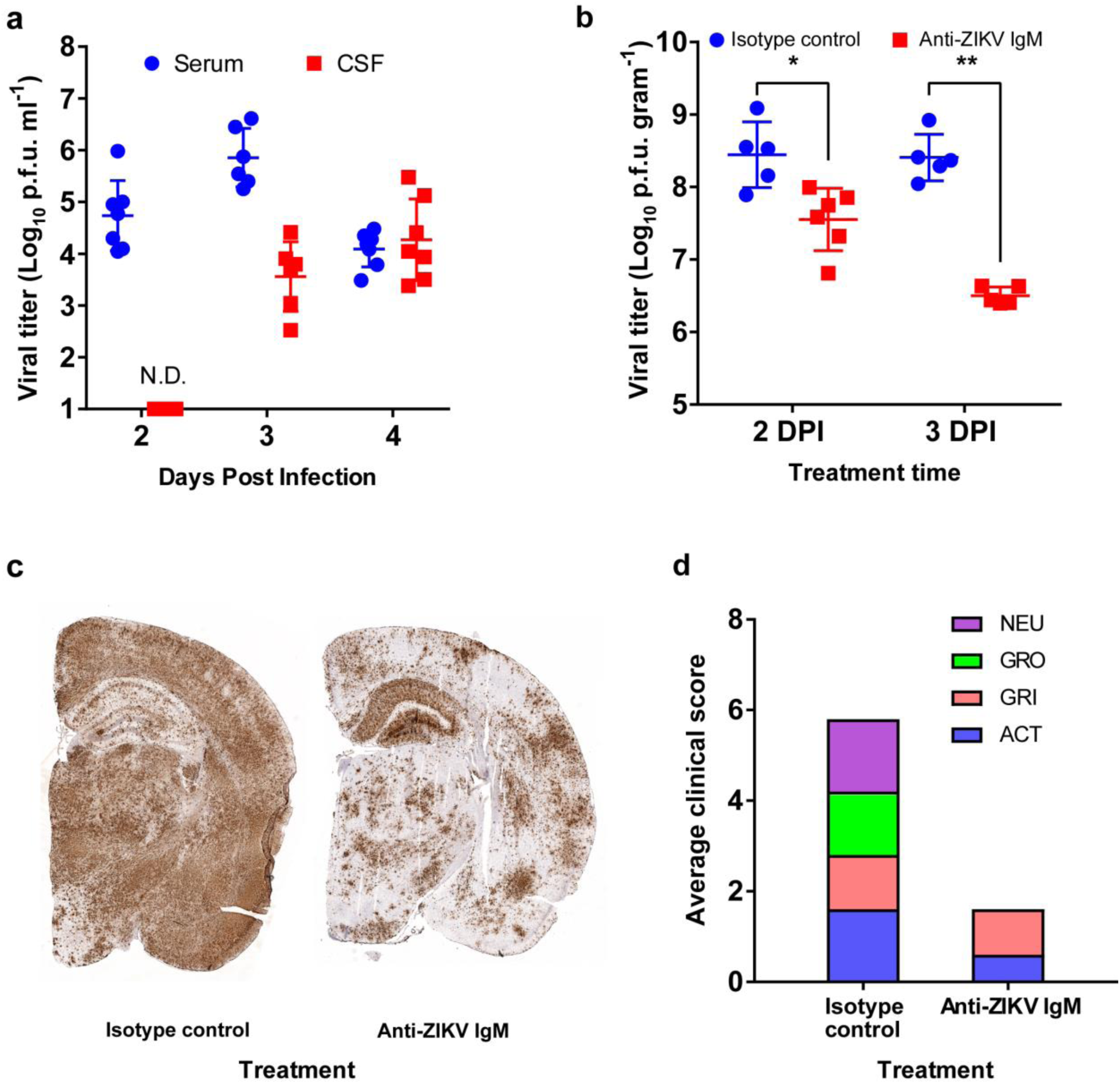
ZIKV in the CSF is critical to brain infection. **a**, IFNAR^-/-^ mice (n= 6–8/time point from two independent experiments) were infected with ZIKV at the footpad as above. The serum and CSF were harvested at 2,3 and 4 DPI as described in the Methods section. The virus titers in the serum (blue circles) and in the CSF (red squares) were measured with a standard focus-forming assay in Vero 76 cells. **b-d**, Anti-ZIKV IgM or isotype control IgM was administered intrathecally to ZIKV-infected mice at 2 or 3 DPI (n= 6 per group per time point from two independent experiments). The brains were harvested at 6 DPI. **b**. The viral loads in the right hemisphere of the brains from the isotype control group (anti-fluorescein IgM, blue circles) and Anti-ZIKV IgM (red squares) were determined. **c**. A representative image of the RNAScope assay for the left hemisphere of the brains (n=4 per group). **d**. Clinical scores (0 to 4, normal to severe) of the tested mice at 6 DPI. NEU: neurological symptoms; GRO: grooming; GRI: grimace scale; Act: Activity.

Our results showed 1) active ZIKV replication in the meninges and the CP and 2) the presence of ZIKV in the CSF prior to active infection in the cortex. Together, these results suggest that ZIKV in the CSF at the early stage of infection is likely the result of infection of the CP and/or the meninges, not a consequence of the infection in the cortex. Hence, we hypothesized that ZIKV circulating in the CSF is responsible for CNS infection. To test this hypothesis, we employed an intrathecal delivery of a ZIKV-neutralizing antibody in our ZIKV mouse model. IFNAR^-/-^ mice were infected with ZIKV subcutaneously in the footpad as described above, and we administered either monoclonal anti-ZIKV IgM antibody (ZKA185 clone(51), mouse IgM isotype) or monoclonal anti-fluorescein IgM antibody (isotype control) intrathecally into the CSF through the cisterna magna at 2 or 3 DPI. The anti-ZIKV IgM antibody showed very strong neutralizing activity against ZIKV in vitro with a PRNT^50^ of 1.3 ng/mL (Supplementary Fig.6). We chose the IgM isotype antibody over the IgG type to maximize availability in the CSF. The clearance rate of IgM in the CSF after an intrathecal delivery is known to be approximately two-fold slower than that of IgG (52). The effect of intrathecal delivery of anti-ZIKV IgM antibody was evaluated by comparing the viral loads in tissues and the clinical symptoms at 6 DPI (Fig. 6b, c, and d). For this model, ZIKV-infected, untreated mice typically die or develop neurological symptoms thus meeting the euthanasia criteria at 7 DPI. Mice treated with anti-ZIKV IgM antibody showed a significantly lower brain viral load and clinical symptoms than those of the isotype control group at 6 DPI. For example, when ZIKV-infected mice were treated with the antibodies at 3 DPI, the mean brain titers of the isotype control and anti-ZIKV IgM treated group were 3.27 x 10^8^ p.f.u./g and 3.28 x 10^6^ p.f.u./g brain tissue, respectively (*p*<0.005, Student’s t-test). We also confirmed such a difference using the RNAScope assay with brain sections (Fig. 6c). Similarly, the group treated with anti-ZIKV IgM showed much milder clinical symptoms compared to those of the isotype control group at 6 DPI (Fig. 6d). Viral loads of peripheral tissues (e.g., serum and spleen) showed no significant or little difference between the groups compared to those of the brain, indicating that the primary effect of antibody treatment was on the infection in the brain (Supplementary Fig. 7a). Intraperitoneal injection of the same antibody did not attenuate viral infection in the brain (Supplementary Fig. 7b). The observed attenuated brain infection by neutralization of the virus in the CSF demonstrated that ZIKV in the CSF at the early stage of infection might be responsible for establishing a lethal infection of the brain.

## Discussion

ZIKV is unique among flaviviruses with regards to its transmission and tissue distribution. Sexual transmission and infection of organs that are separated from the circulation (e.g., eyes, brain, fetus, and reproductive tract(7,53–55)) establish ZIKV as a unique flavivirus that can cross various biological barriers. While nonneurological ZIKV infections cause no to mild symptoms in humans, ZIKV infection in the brain causes much more severe clinical manifestations, such as encephalitis, microcephaly, and other ZIKV-associated neurological symptoms. This fact underscores the importance of defining the mechanism by which ZIKV crosses brain barriers.

Here, we show that ZIKV might invade the brain from the circulation by exploiting the B-CSF barrier instead of the BBB. The BBB is the primary barrier that prevents most infectious agents from entering the brain. While many neurotropic viruses such as VEEV(23) are known to use the BBB as an entry route to the brain, the mechanism by which ZIKV enters the brain remains unclear. Some experimental findings highlighted the involvement of the breakdown of the BBB by ZIKV-infected astrocytes(56); however, this may be a result of the brain infection rather than the brain entry mechanism for the following reasons: 1) later onset of BBB destruction than the initiation of brain infection (6 DPI compared to 3 DPI) and 2) the undefined origin of the virus that infected the astrocytes.

Our understanding of the role of CP in the context of brain infection is very limited; however, the CP exhibits many biological characteristics that can contribute to the entry of neurotropic viruses into the brain. Unlike the endothelial cells of the BBB, the CP endothelial cells are fenestrated and leaky (i.e., fenestrated blood capillaries with diaphragmed fenestrae) with fenestrae ranging between 60 and 80 nm in size(57). Although some reports showed that the maximum size of molecules that can pass through the fenestrae may be smaller than the physical size of the fenestrae(58), the fenestrated endothelial cells of the CP could provide an ideal milieu for viruses to spread out of the blood and into the cisternae of the CP.

Pericytes are in direct contact with endothelial cells; hence, viruses that have come out of the circulation through endothelial cells might directly contact nearby pericytes. Recent studies, which used fluorescent proteins under control of the NG2 (Cspg4) and PDGFRB promoters and single-cell sequencing of mouse brain cells, have identified several pericyte subpopulations in the brain, indicating a diversity of pericytes in both function and differentiation(59, 60). In addition, pericytes in the brain are known to differentiate into other cell types, such as fibroblasts, indicating a substantial plasticity of pericytes(26, 27). While brain perivascular pericytes have been extensively studied at the BBB level(61, 62), little is known about the biological functions of CP pericytes, especially the biological roles of CP pericytes with respect to viral invasion through the B-CSF barrier. Here, our study presents a novel finding that CP pericytes can be infected with ZIKV and might provide a local amplification site within the CP. Considering the variety of cytokines that are known to be secreted from pericytes(39, 63), it would be important to understand other roles of ZIKV-infected CP pericytes in the viral invasion of the brain as well as in local inflammatory signals. The role of Axl in ZIKV infection has been studied extensively in mechanisms pertaining to: 1) cellular entry receptor and 2) attenuation of innate immune responses. ZIKV is known to bind to TAM receptors by using the ligands as a bridge, whereby TAM ligands (e.g., Gas6 or protein S) bind to phosphatidylserine exposed on virus particles and bridge virions to TAM receptors and utilizes the receptors as cellular receptors for entry, as well as attenuators of innate immune responses in the infected cells (i.e., the viral apoptotic mimicry mechanism)(46,47,64,65). However, some evidence indicates that the dependency of ZIKV on TAM receptors for infection might be cell-type or model specific (e.g., IFN dependency) (48, 49). Our data clearly demonstrate that ALX is important for ZIKV infection in HBVP. A future study to understand to what extent the infection of pericytes is mediated via TAM receptors will be of great interest.

With respect to the blood-brain interface, CP epithelial cells serve as a selective barrier and a professional transporter between the blood and the CSF(28). The tight junctions between CP epithelial cells (e.g., zonula occludens) prevent paracellular diffusion. CP epithelial cells transport biological molecules across the epithelial barrier via transcytosis, which can be exploited by neurotropic viruses as a brain entry mechanism(30,66,67). A very limited number of studies have been conducted with CP epithelial cells with respect to neuroinvasion. John Cunningham virus and Chikungunya virus are known to infect CP epithelial cells in vitro(37, 68); however, to our knowledge, our study shows for the first time neurotropic viruses traversing the CP epithelial cell layers without active replication. Importantly, interferon gamma (IFN-γ), the Th2-type cytokine interleukin-4 (IL-4), and MPC-1 are known to enhance transcytosis of IgA and HIV(69, 70). Confirmation of the release of these cytokines by ZIKV-infected pericytes and enhancement of the transcytosis of ZIKV would explain how pericytes contribute to ZIKV brain entry via the B-CSF barrier, which was demonstrated as enhanced transport of ZIKV through the cells in our coculture model (Fig. 5).

Overall, our study demonstrates a novel CNS invasion mechanism exploiting pericytes around fenestrated capillaries and epithelial cells in the B-CSF barrier of the CP by ZIKV. The ensemble of fenestrated capillaries, pericytes, and epithelial cell layer serves as a barrier and transporter in other organs, such as the eye and the reproductive tract. Hence, our study might also shed light on the unique secretion and invasion profiles of ZIKV and new approaches to prevent the transmission of the virus.

## METHODS

### Viruses

The ZIKV strains MR766 and PRVABC59 (GenBank: KX087101) were provided by Dr. Barbara Johnson at the CDC. The following reagent was obtained through BEI Resources, NIAID, NIH: ZIKV strains PLCal_ZV (Human/2013/Thailand; GenBank: KF993678) and DAK AR 41524 (Mosquito/1984/Senegal; GenBank: KX198134.2). ZIKV was amplified in Vero 76 cells (ATCC CRL-1587), and the titer was determined in Vero 76 cells with a standard methylcellulose overlay method described in a previously published paper(71). VEEV TC-83 was obtained from USAMRIID (a gift from Dr. Connie Schmaljohn) and amplified in BHK-21 (ATCC CCL-10).

### Antibodies

Mouse-4G2 (monoclonal antibody against anti-flavivirus group antigen, clone D1-4G2-4-15, ATCC), Hu-4G2 (recombinant monoclonal antibody of clone D1-4G2-4-15 with human IgG Fc produced in Nicotiana tabacum, a gift from Dr. Nobuyuki Matoba), anti-TTR (Abcam), anti-CD31 (Invitrogen), anti-PDGF receptor beta (Abcam), anti-alpha smooth actin (Abcam), anti-Axl (R&D systems), FITC-isolectin B4 (Sigma Aldrich), Alexa Fluor 647-conjugated goat anti-rabbit IgG (Jackson ImmunoResearch), Alexa Fluor 549-conjugated goat anti-human IgG (Jackson ImmunoResearch), and Alexa Fluor 488-conjugated goat anti-rat IgG (H+L) (ThermoFisher) were used for the immunofluorescence assay. All antibodies were validated for their specificity using mock or isotype antibody controls.

Mouse IFNAR-1 neutralizing antibody (clone MAR 5A3) and isotype control (clone MOPC-21) were purchased from BioXcell. Anti-Axl (AF154), anti-GAS6 (AB885), and isotype control IgG (AB-108-C) were purchased from R&D systems. The human type 1 IFN neutralizing antibody mixture was purchased from PBL Assay Science. Anti-Zika mouse IgM antibody (Clone ZKA185) and an isotype control mouse IgM antibody (anti-fluorescein, clone 4-4-20) were obtained from Absolute Antibody.

### Cells

Vero 76 cells (ATCC 1587) were obtained from ATCC and maintained in MEM-E with 10% FBS and L-glutamine. Primary human brain vascular pericytes (HBVPs) were obtained from ScienCell (Cat No. 1200) and maintained using the manufacturer’s protocol and their pericyte culture media with 2% FBS, 1 x pericyte growth supplement, and 1 x penicillin and streptomycin.

Mouse choroid plexus pericytes (MCPPs) were isolated according to the protocol described by Boroujerdi et al.(72) with modifications. Choroid plexuses isolated from 4-5 mice at 5 weeks of IFNAR^+/-^ were incubated in 1 mL of a papain/DNase I digestion solution (20 units of papain, 1 mM L-cysteine, and 0.5 mM EDTA in HBSS) at 37 °C for 50 min. After trituration of the choroid plexuses by 7 passages through a 19-gauge, followed by 7 passages through 21-gauges, the cell mixture was mixed with the same volume of 22% BSA to neutralize the papain. Cells were harvested by centrifugation and suspended in 3 mL of endothelial cell growth medium (F12 medium with 10% FBS, 1x penicillin and streptomycin, 30 µg/mL endothelial cell growth supplement (Millipore), 2.5 µg/mL ascorbate, 1 x GlutaMax, and 40 µg/mL heparin). Cells were plated on collagen type I-coated 35 mm dishes, and unattached cells were removed 2 hours after seeding. Pericytes were selected and cultured in pericyte growth medium (Pericytes Medium-mouse, Sciencell) for four passages prior to experiments. HIBCPP cells were obtained from Dr. Christian Schwerk’s lab (Univ. of Heidelberg) and maintained according to a method published previously(50, 73).

### Mouse models and virus infection

Heterozygous of Ifnar1tm1.2Ees mice (IFNAR^-/-^) (interferon ɑ/β receptor deficient on the C57BL/6J background) were purchased from Jackson Laboratory (#028288). AG129 mice (interferon ɑ/β/ɣ receptor deficient on the S129 background) were purchased from Marshall Bioresources. Mice were subsequently bred and housed at the University of Louisville. The genotypes of test mice were determined following the provider’s protocol. Mice of both sexes between 4 and 6 weeks old were used for all experiments. The animal experiments were carried out in strict accordance with the recommendations in the Guide for the Care and Use of Laboratory Animals of the National Institutes of Health. The animal experiments were conducted following an approved protocol (Protocol Number: 16533) by the University of Louisville Institutional Animal Care and Use Committee. Four-to six-week-old mice of both sexes were anesthetized under isoflurane and infected subcutaneously with 1000 p.f.u of virus (100% lethal in IFNAR^-/-^ or in AG129 mice) diluted in PBS at the rear foot pad.

### In situ chromogenic RNA hybridization (i.e., RNAScope assay)

Mice were euthanized at the time points denoted in the experiments. The brains were harvested, and the left hemisphere of each brain was immediately placed in 10% buffered formalin and fixed for 24 to 48 hours followed by washing with 70% ethanol. The fixed brains were embedded in paraffin and sliced at a thickness of 5 µm and mounted on glass slides. Target RNA in the tissue sections was detected using RNAScope® 2.5 Assay (Advanced Cell Diagnostics) and a specific probe according to manufacturer’s protocol with a modification of 25 min of protease treatment instead of 30 min(74). Stained tissues were counterstained with Gill’s hematoxylin (American Master Tech Scientific), dehydrated, and mounted with glass coverslips using Cytoseal XYL (Fisher Scientific). Tissues were then visualized with a bright-field microscope. The image background was corrected with ImageJ as described elsewhere. (http://imagejdocu.tudor.lu/doku.php?id=howto:working:how_to_correct_background_illumination_in_brightfield_microscopy). High-resolution cross-section images were created using the Hugin Panorama photo stitching program (hugin.sourceforge.net) with small-field images for each image taken with a 10 × objective.

### Determination of viral loads in the brain

Mice were infected subcutaneously with ZIKV strain PLCal_ZV. At 4 DPI, the mice were euthanized and transcardially perfused with 15 mL of PBS prior to the removal of the brains. The cerebral surface vasculature and associated meninges were harvested from the ventral and dorsal sides of the brains according to a protocol published by others(75). Choroid plexus was isolated from all of the brain ventricles and pooled. Approximately 100 mg of cerebral cortex was harvested from the isocortex. Each tissue was homogenized in 0.5 mL of RNAzol RT (Molecular Research Center, Inc), and total RNA was isolated according to the manufacturer’s protocol. The total RNA was subjected to reverse transcription with 200 U of Maxima H Minus Reverse Transcriptase and 2 µM of random hexamer in a total volume of 25 µL. qRT-PCR was performed in a multiplex format with 500 nM of ZIKV-specific primers and 250 nM of probes and a GAPDH-specific primer/probe set (4352339E, Applied Biosystems). The following ZIKV primer/probe sequences: Zika 1087 NW_fwd 5’-CCGCTGCCCAACACAAG-3’ and Zika 1087 NW_rev 5’-CCACTAACGTTCTTTTGCAGACAT-3’, and 5’/6-FAM/AGCCTACCTT/ZEN/GACAAGCAATCAGACACTCAA/IABkFQ/3’ were modified specific to the PLCal_ZV strain based on the sequences reported by Lanciotti et al.(76).

### Immunostaining of whole-mount choroid plexus

Mice infected with 1000 p.f.u of ZIKV as described above were euthanized, and the brains were harvested immediately. The brains were rinsed twice in ice-cold HBSS, and the choroid plexuses were isolated from all ventricles (lateral, third, and fourth) under a stereoscope. The isolated choroid plexuses were incubated in HBSS with Mg^2+^ and Ca^2+^ containing 100 µg/mL DNase I (Akron Biotechnology) for 20 min at 37 °C followed by washing in PBS twice. The choroid plexuses were fixed with 4% paraformaldehyde in PBS for 20 min at 4 °C and washed twice in PBS with 0.1% Triton X-100. The fixed choroid plexuses were blocked with a blocking-permeabilization buffer (20% horse serum and 0.3% Triton-X100 in PBS) for two hours at 4 °C. For imaging, the fixed choroid plexuses were incubated overnight with antibodies for specific cell markers: anti-CD31 for endothelial cells, anti-ZIKV E for ZIKV, and anti-PDGF receptor beta for pericytes, diluted with an antibody dilution buffer (2% goat serum and 0.1% Triton X-100 in PBS). Next, after two washes, the tissues were incubated with fluorophore-tagged secondary antibodies. For endothelial cell staining with FITC-isolectin B4, 10 µg/mL FITC-isolectin B4 (Sigma Aldrich) was added to the secondary antibody solution. These tissues were washed and further stained with Hoechst 33342 at a concentration of 1 µg/mL for 20 min. After mounting with ProLong Diamond antifade mount, immunofluorescent images were acquired using a Zeiss LSM 710Duo/Live5 confocal laser scanning fluorescence microscope. Images were processed using Imaris (Oxford Instruments) or FIJI (https://imagej.net/Fiji) software.

### Collection of CSF and serum

CSF was collected according to the protocol by Nastasia et al.(77). In brief, mice were anesthetized with an intraperitoneal injection of tribromoethanol (250 mg/kg), and the scalp was shaved. The dura of the cisterna magna was exposed by an incision under a stereoscope. CSF (1∼5 µL per mouse) was harvested from the cisterna magna by a puncture of the dura with a capillary pipette (with an inner diameter of 50 µm). Blood contamination was determined by visual inspection of red blood cells in the CSF samples. CSF with blood contamination was excluded from further assays. After CSF collection, the blood was collected from the heart. The collected blood was incubated at room temperature for 20 min to allow clotting. The serum was harvested after centrifugation at 8,000 x g at 4 °C. Blood-free CSF was suspended in 100 µL of complete media (MEM-E, 10% FBS, and 25 mM HEPES) and clarified by low-speed centrifugation for 5 min at room temperature.

Collected samples were stored at −80 °C until use. Viral loads in the CSF and the serum were determined by following a standard methylcellulose overlay method as described previously(71).

### Virus infection of primary HBVP and MCPP cells

Cells were seeded at a density of 50,000 cells/ well in 24-well plates coated with poly-L-lysine (for HBVPs) or collagen type-1 (for MCPPs) and incubated overnight prior to infection. Cells were infected with 10,000 p.f.u./well (∼0.2 MOI) at 37 °C for one hour and washed with PBS twice to remove unbound virus. Cells were replenished with fresh media with anti-IFN neutralizing antibodies and incubated at 37 °C with 5% CO^2^. At 1, 3, and 5 DPI, the cell culture supernatants were harvested, and the viral titers in the supernatants were determined in Vero 76 cells by following a standard methylcellulose overlay method as described above.

### Immunostaining of HBVP

HBVP cells were seeded as described above and infected with ZIKV at a m.o.i. of 1 at 37 °C for 4 hours. For imaging cells were fixed in ice-cold methanol at room temperature for two minutes, then blocked with 5% normal goat serum and 0.1% Triton X-100 in PBS for one hour. Cells were then incubated at 4 °C overnight with primary antibodies for specific markers followed by fluorophore-conjugated secondary antibodies. All antibodies were diluted 1:400 in 1% BSA, 0.05% saponin in PBS. The glass slips were washed in PBS and further stained with Hoechst 33342 at a concentration of 1 µg/mL for 20 min. After mounting with ProLong Diamond antifade mount, immunofluorescent images were acquired using a Zeiss LSM 710Duo/Live5 confocal laser scanning fluorescence microscope. The proportion of infected cells were determined by image analysis. For each image, 16 field-of-view taken with a 10 X objective were combined together and numbers of nuclei (i.e., total cell numbers) and ZIKV-E positive cells in each image were determined with Imaris software.

### ZIKV-positive cell counting with flow cytometry

HBVP cells were seeded into T-25 cell culture flasks at a density of 600,000 cells/flask and were incubated for 2 days at 37 °C and 5% CO^2^ prior to infection. The cells were then infected with ZIKV (0.1 MOI) by incubating with virus at 37 °C for 4 hours. After incubation, the cell supernatant was aspirated and replaced with fresh media. At the indicated timepoints (1, 2, 3, 4, or 5 days), the cells were detached from the culture surface by trypsinization with TrypLE Express (Gibco). After neutralization and trituration with pericyte culture media, the cells were passed through a 70 µm nylon filter (VWR International), washed twice with 1x PBS, and passed through a 70 µm filter again to ensure a single cell suspension. The cells were then fixed for 30 minutes at 4 °C in 4% paraformaldehyde and 0.1% saponin in 1x PBS. After washing of the cells with wash buffer (0.2% BSA, 0.1% saponin in 1x PBS) twice, the fixed cells were then stained with mouse-4G2 antibody (0.5 µg/mL in PBS with 1% BSA, 0.1% saponin) at room temperature for 1 hour. After incubation, the cells were washed with wash buffer once and then incubated with 1.5 µg/mL of Alexa Fluor 488-conjugated anti-mouse IgG in PBS with 1% BSA, 0.1% saponin for 1 hour at room temperature. Finally, the HBVP cells were washed three times with wash buffer and subjected to flow cytometry analysis. Flow cytometry analysis was performed using a BD LSRFortessa (BD Bioscience) flow cytometer with BD FACSDiva software (BD Bioscience). Forward scatter and side scatter voltages were optimized to place the population of interest on scale using an uninfected control sample. A histogram was created with the fluorescent intensity and cell counts. Cells with significantly higher fluorescent intensity than mock infected cells were considered ZIKV-positive cells. Further analysis was performed using FlowJo software (FlowJo).

### AXL and GAS6 blockade

HBVP grown in 24-well plates were preincubated with 15 μg/ml anti-AXL antibody, 5 μg/ml anti-GAS6 antibody, or goat polyclonal IgG control (n=4 per group) for 3 hrs at 37°C prior to infection. Cells were infected with ZIKV at an M.O.I of 1 for 4 hours and washed with PBS once. Cells were replenished with fresh media with the half concentration of antibodies and were incubated for two days. The total RNA was isolated from the cells and subjected to qRT-PCR for viral RNA quantitation. Viral titers in the supernatants were enumerated in a plaque assay as described above.

### Neutralization of ZIKV in the CSF

Mice (IFNAR^-/-^, n=6 per group from three independent experiments) were infected with 1000 p.f.u. of ZIKV subcutaneously at the footpad. At 2 DPI, the mice were treated with anti-fluorescein mouse IgM (the isotype control group) or ZKA-185 mouse IgM (ZIKV neutralizing antibody group) via an intrathecal administration. For intrathecal administration, mice were anesthetized with an i.p. injection of a ketamine/xylazine mixture (100 and 5 mg/kg, respectively), and isoflurane was provided during the surgery. After the dura of the cisterna magna was exposed as described above in the “Collection of CSF and serum” section, 3 μL of antibody (1 mg/mL in PBS, 3 µg/mouse) was slowly administered into the cisterna magna cistern (3 µL/5 min) using a capillary pipette (tip size of 30 µm) controlled with a pneumatic microinjector (World Precision Instrument). After two min of resting after the capillary was retracted from the puncture, the incision was sutured, and the animals were returned to the cages. At 8 DPI, the animals were euthanized followed by cardiac perfusion with 15 mL of PBS. The right hemispheres of each brain were immediately placed in 10% buffered formalin and fixed for 24 to 48 hours for use in a ZIKV-RNAScope assay as described above. The remaining left hemispheres were homogenized in VIM at 10% (weight/volume) and then cleared by centrifugation. The virus titers and viral RNA copy numbers in the 10% brain homogenate were determined by the methylcellulose overlay method and qRT-PCR with a standard curve method.

### Electron micrograph

IFNAR^-/-^ mice were infected with ZIKV as above. At 4 DPI, mice were anesthetized with an intraperitoneal injection of tribromoethanol (250 mg/kg) and then transcardially perfused with 20 mL of ice-cold phosphate buffer saline to wash out the residual blood from the cerebral circulation. After perfusion with cold PBS, mice were perfused with 15 mL of fixative (2% paraformaldehyde, 2% glutaraldehyde in 0.1 M phosphate buffer, pH 7.4). The brains were harvested and then kept in the fixative. The choroid plexuses were isolated under a stereoscope and then fixed further at 4 °C for three days. After washing with phosphate buffer, the tissues were treated with 1% osmium tetroxide for two hours and then embedded with Durcupan resin. Sections were imaged with a transmission electron microscope (Hitachi HT7700).

### Virus transport through Transwell inserts

To establish barriers on Transwell plates, HIBCPP cells were seeded in the upper compartment of 24-well format Transwell inserts with a pore size of 0.4 µm and a density of 2 x 10^6^/cm^2^. After two weeks of culture, the transepithelial electric resistance (TEER) of each insert was confirmed to be > 600 Ω/insert. Then, cells were further cultured in reduced FBS HIBCPP media with 2% FBS instead of 10% for one day and then used for experiments. HBVP cells grown in 24-well plates overnight were infected with the ZIKV strain PLCAL_ZV at an MOI of 0.1 for 4 hours and then washed once with PBS before fresh culture media was replenished. At 2 DPI, Transwell inserts with fully developed HIBCPP barriers were placed in the well. After 24 hours of coculture, the cell supernatants from the upper and lower chambers were harvested, and the viral titers were measured as described above. At 5 DPI, the integrity of the barrier functions was tested with HRPO-conjugated antibody diffusion assay(78).

## ACKNOWLEDGMENTS

We thank BEI Resource and Dr. Barbara at the CDC for providing ZIKV strains. We also thank Dr. Christian Schwerk for providing the HIBCPP cell line. We are grateful to Robert Adcock, University of Louisville Center for Predictive Medicine Animal Core, University of Louisville EM Core, and Iowa State University Comparative Pathology lab for helpful technical support. We also thank Dr. David Magnuson for helpful comments and insights and Christine Yarberry, Gretchen Holz, and Dr. Yong-kyu Chu for establishing the assays. This research was supported by the Collaborative Matching Grant Program from School of Medicine, University of Louisville. All experiments involving animals and animal tissues were performed in accordance with protocols approved by the University of Louisville Institutional Animal Care and Use Committee.

---

**Supplementary Figure 1.**
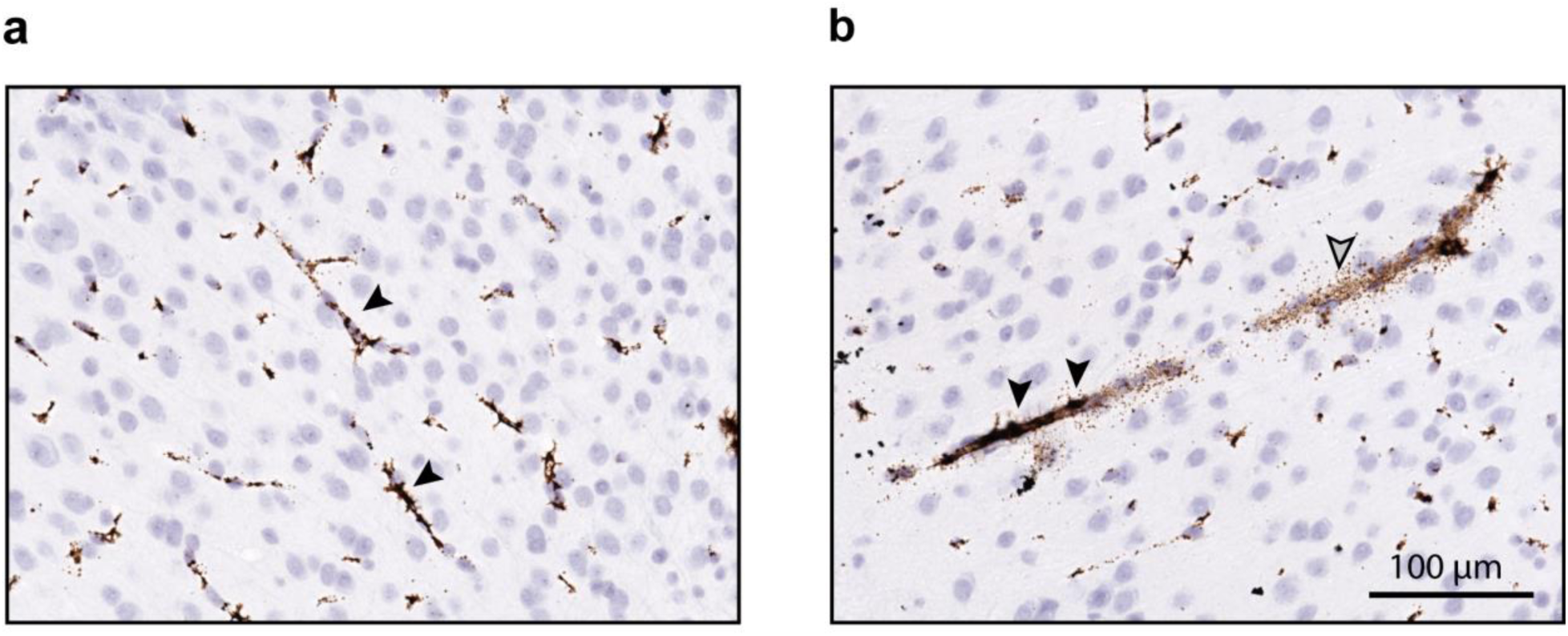
VEEV TC-83 infects the brain at the level of BBB. The brains infected with Venezuelan equine encephalitis virus (strain TC-83) showed widely distributed strong positive staining around the capillaries in the cortex. Representative images of viral RNA staining with RNAScope assay of the brain cortex of TC-83 infected AG129 mice (n=6 from two independent experiments). Images were taken with a 40 x objective. Cortex capillaries showed strong positive straining (dark brown) for viral RNA. Black and gray arrowheads indicate virus-infected cells and virus -specific staining, respectively.

**Supplementary Figure 2.**
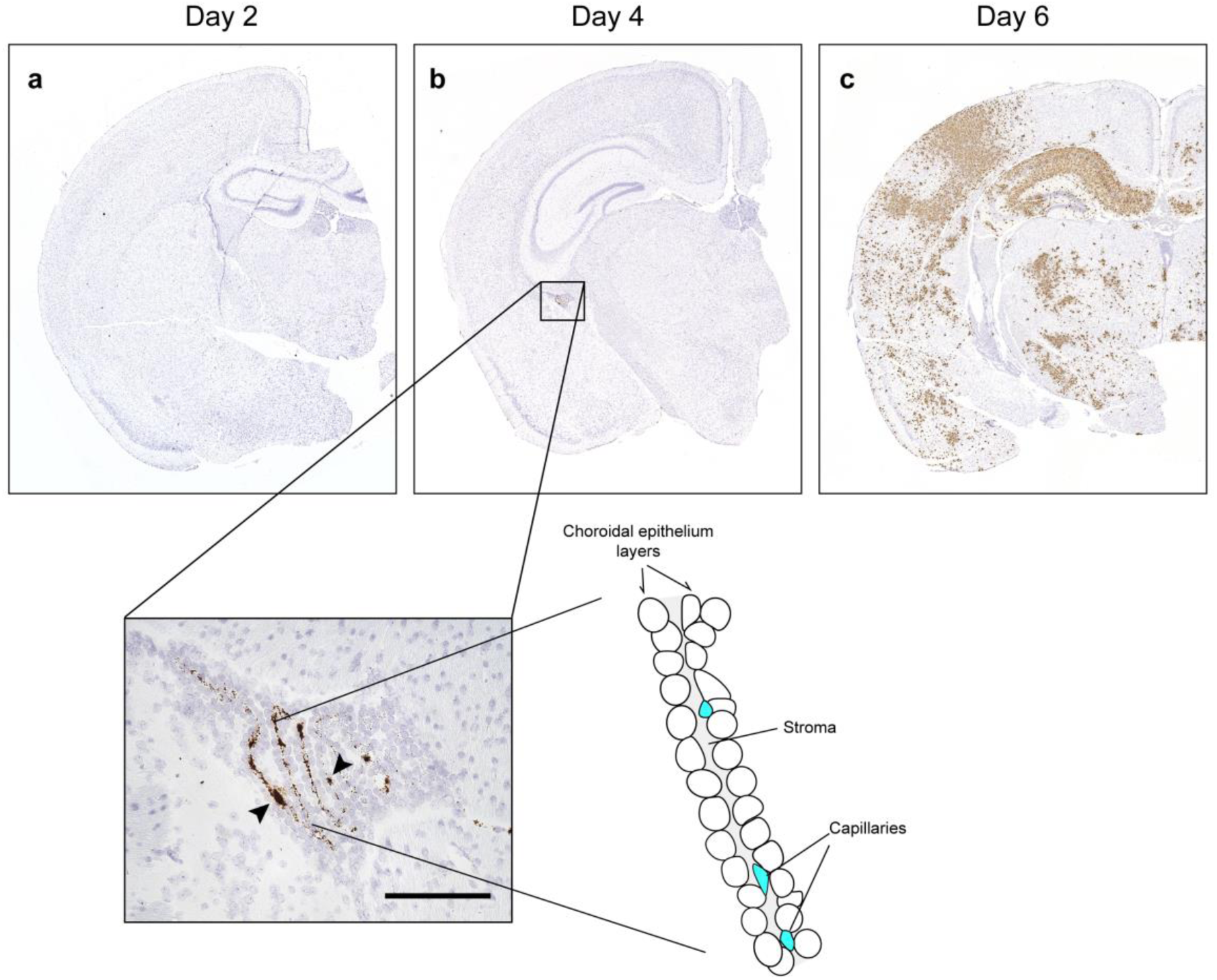
ZIKV appears the stroma area of the choroid plexus first in the brain of the AG129 mice infected with ZIKV. Brains from AG129 mice infected with PLCal_ZV (n=3/timepoint, 1000 puf/mouse) were subjected to RNAScope assay to detect viral RNA. For **a**-**c**, representative images were shown from brains harvested at 2, 4, and 6 days post infection. Black arrowheads indicate virus-infected cells. Scale bars, 100 µm.

**Supplementary Figure 3.**
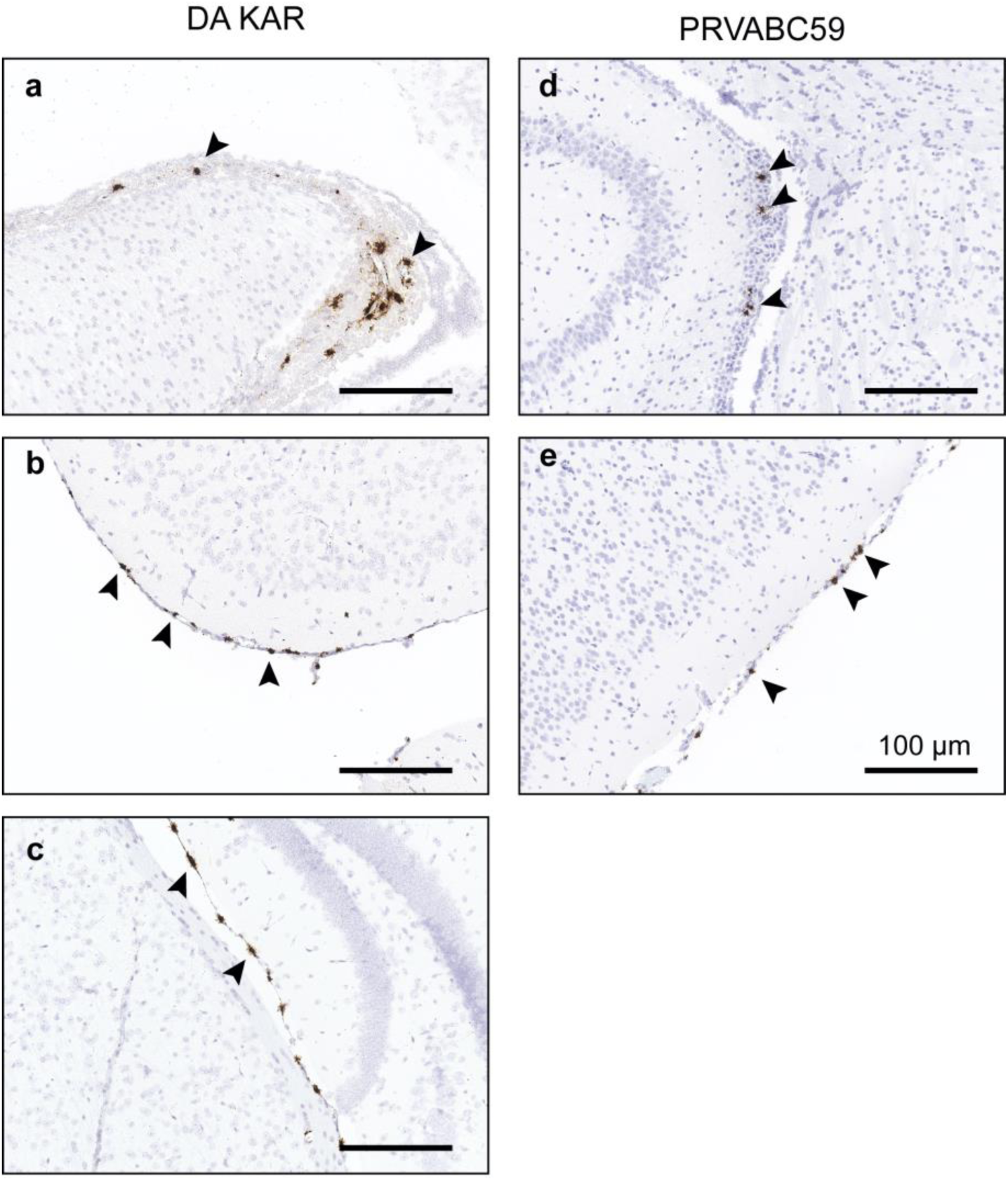
Infection of the choroid plexus and meninges of the brain at the early stage of infection is a common feature of ZIKV brain infection. Representative image of the brains of AG129 mice infected with ZIKV strain DA KAR (**a**-**c**) and PRVABC59 (d-e) (n=3 per group). The brains were harvested at 3 d.p.i. and were analyzed with RNAScope assay with a specific probe against the ZIKV.

**Supplementary Figure 4.**
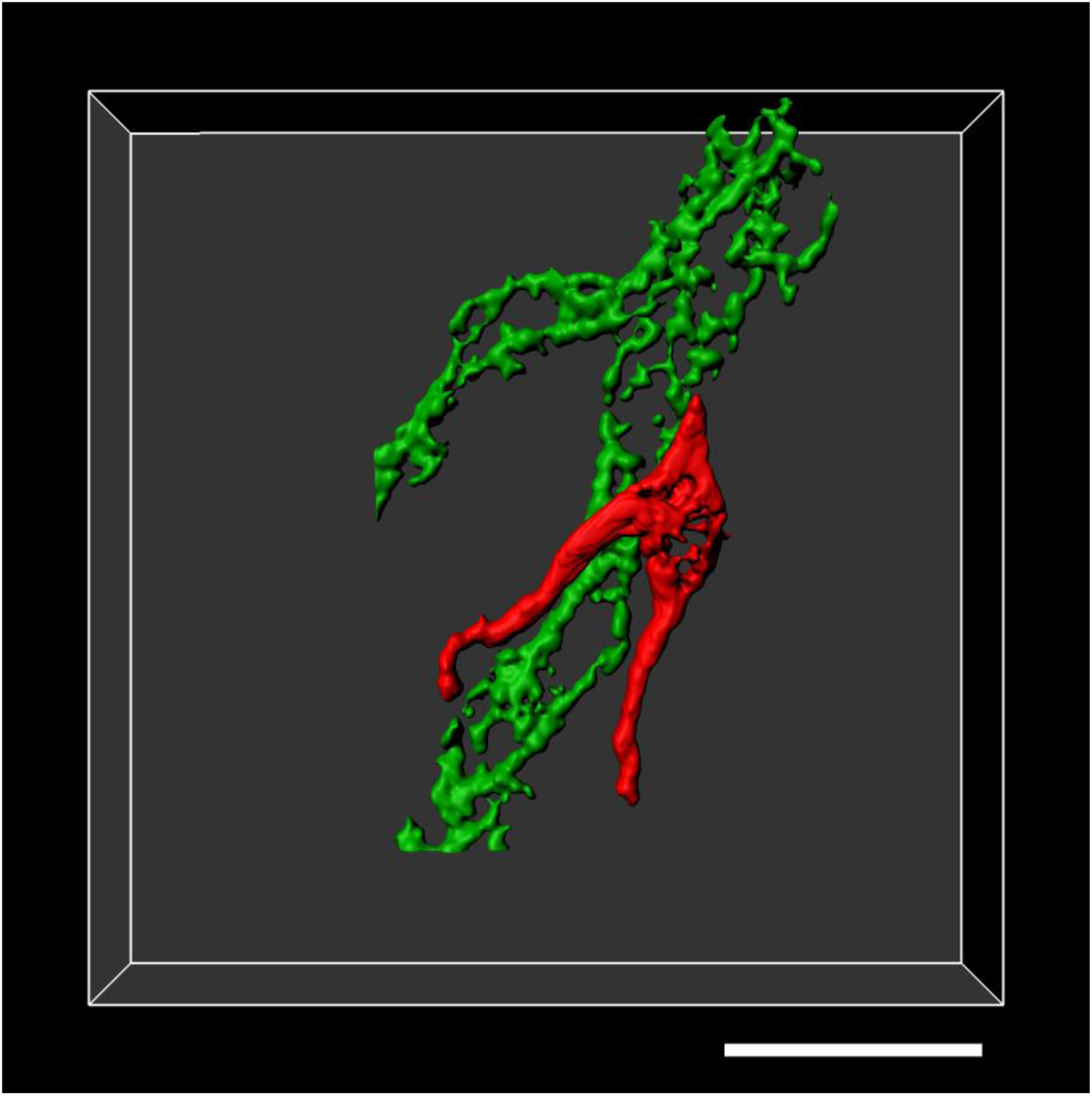
A reconstituted 3D model of a ZIKV-infected pericyte. A reconstructed 3-D image of Fig. 3b. Images were acquired with a confocal laser scanning microscopy and a three-dimensional image was reconstructed with image analysis software Imaris using its surface modeling function. CD31 (green) and ZIKV-E (red) were used as markers for endothelial cells and ZIKV-infected cells. Scare bar, 20µm.

**Supplementary Figure 5.**
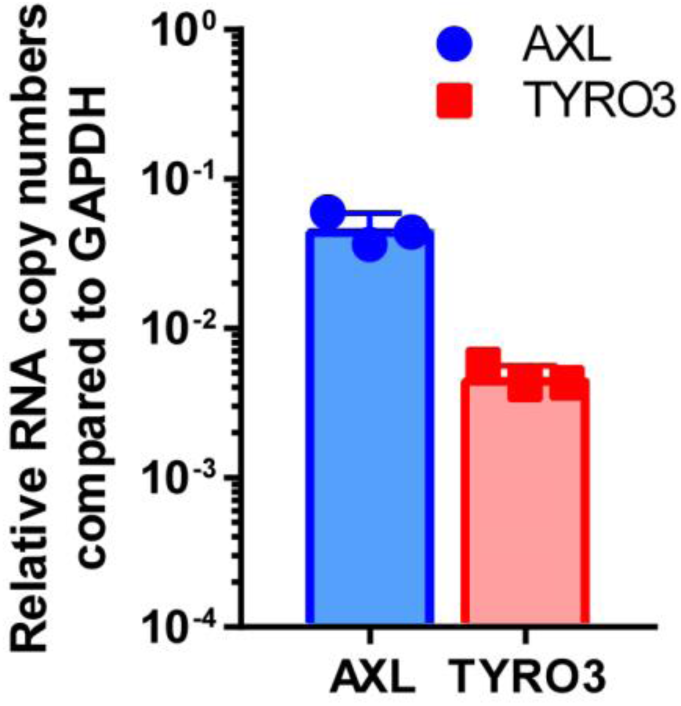
AXL expression in HBVP cells. The total RNA was isolated from HBVP cells (n=3) grown in a 24-well plates for two days and the expression of AXL and TYRO3 in HBVP were determined with gene-specific qRT-PCR assays (TaqMan Gene Expression Assay ID: Hs01064444_m1 and Hs03986773_m1 for AXL and TYRO3, respectively) and normalized to the expression of GAPDH mRNA in each sample.

**Supplementary Figure 6.**
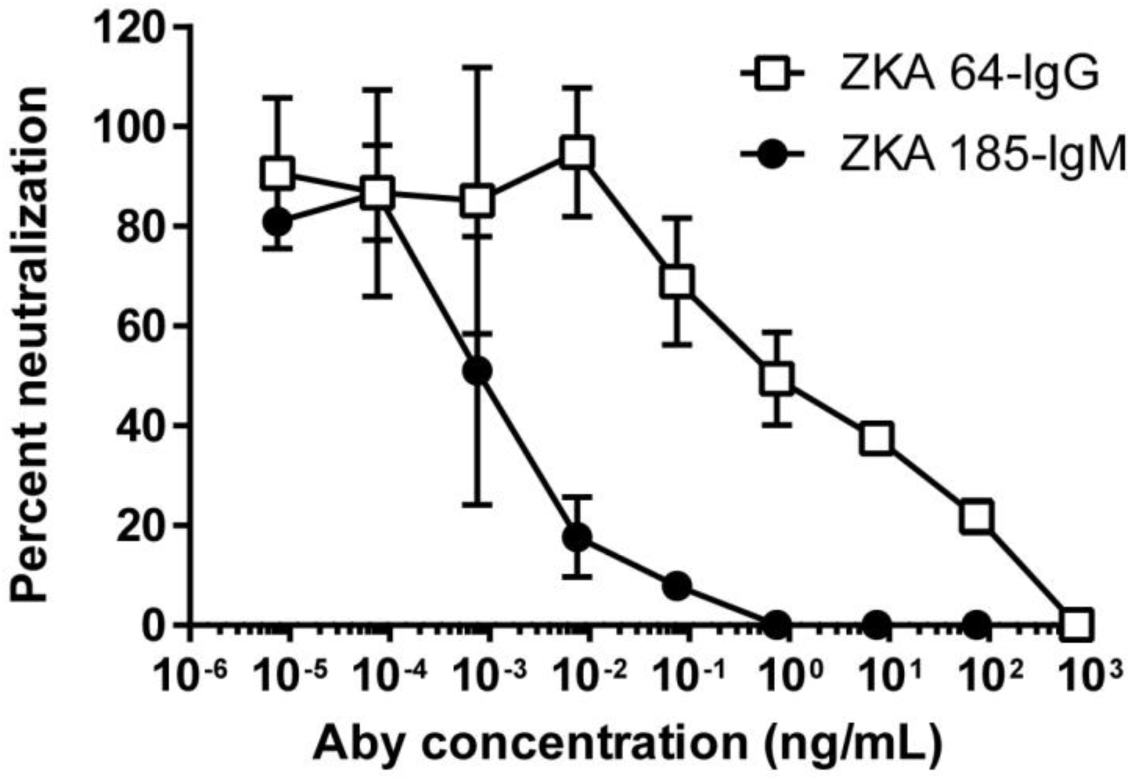
Plaque Reduction Neutralization Activity of antibody used for in vivo neutralization. ZIKV-specific antibodies, clone ZKA 64 and ZKA 185 were serially diluted in cell growth media with HEPES (12.5 mM) and incubated with ZIKV strain PLCal_ZV (100 p.f.u./sample) for one hour at 37 °C. Vero 76 cells grown overnight in 12-well plates were infected with the antibody-virus mix and 5 days later viral plaques were developed by crystal violet staining. Anti-fluorescein mouse IgG and anti-fluorescein IgM were used as non-neutralizing antibody control (10 ng/mL)

**Supplementary Figure 7.**
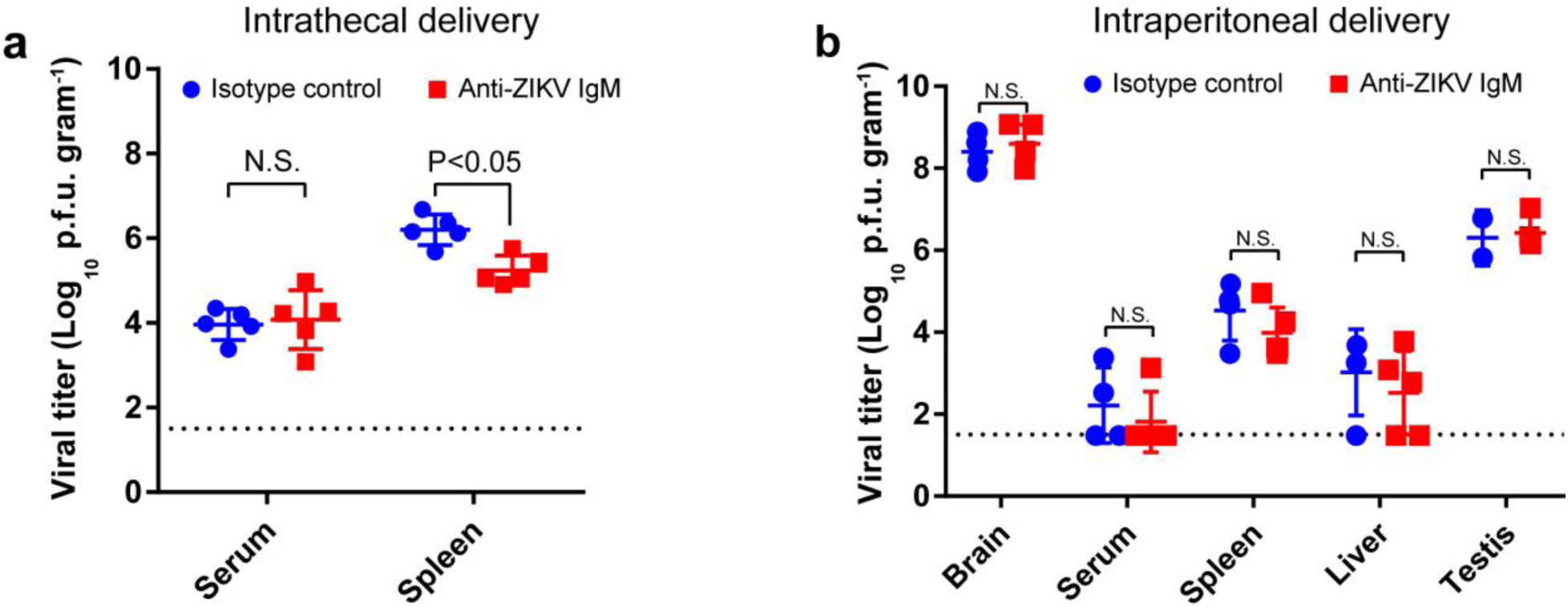
In vivo effect of ZIKV neutralizing antibody delivered via the intrathecal or intraperitoneal routes. **a**, Intrathecal delivery of neutralizing antibody did not affect viral growth in peripheral tissues as much in the brain. Viral loads of serum and the spleens of ZIKV-infected mice treated either with isotype control (blue circles) or with ant-ZIKV IgM showed no significant (serum) or less extent difference (spleen) than for the brains (6 d.p.i, n= 5-6/group). **b**, Intraperitoneal delivery of neutralizing antibody did not show any difference in viral replication in tissues, including brain. Antibodies (n=4-5/group, 3 µg/mouse which is the same dose used for intrathecal delivery) were administrated intraperitoneally at 3 d.p.i. and the mice were euthanized at 7 d.p.i. Viral loads were determined with 10 % tissue homogenates. N.S. no significance by Student t-test *P*> 0.05.

